# Role and Dynamics of Vacuolar pH during Cell-in-Cell mediated Death

**DOI:** 10.1101/2020.11.03.364877

**Authors:** Yan Su, He Ren, Meng Tang, You Zheng, Bo Zhang, Chenxi Wang, Xinyu Hou, Zubiao Niu, Lihua Gao, Zhaolie Chen, Tianzhi Luo, Qiang Sun

## Abstract

The non-autonomous cell death by entosis was mediated by the so-called cell-in-cell structures, which were believed to kill the internalized cells by a mechanism dependent on acidified lysosomes. However, the precise values and roles of pH critical for the death of the internalized cells remained undetermined yet. We creatively employed keima, a fluorescent protein that displays different excitation spectra in responding to pH changes, to monitor the pH dynamics of the entotic vacuoles during cell-in-cell mediated death. We found that different cells varied in their basal intracellular pH, and the pH was relatively stable for entotic vacuoles containing live cells, but sharply dropped to a narrow range along with the inner cell death. In contrast, the lipidation of entotic vacuoles by LC3 displayed previously underappreciated complex patterns associated with entotic and apoptotic death, respectively. The pH decline seemed to play distinct roles in the two types of inner cell deaths, where apoptosis is preceded with moderate pH decline while a profound pH decline is likely to be determinate for entotic death. Whereas the cancer cells seemed to be lesser tolerant to acidified environments than non-cancerous cells, manipulating vacuolar pH could effectively control inner cell fates and switch the ways whereby inner cell die. Together, this study demonstrated for the first time the pH dynamics of entotic vacuoles that dictate the fates of internalized cells, providing a rationale for tuning cellular pH as a potential way to treat cell-in-cell associated diseases such as cancer.

## Introduction

Cell-in-cell structure (CICs) refers to a unique type of cellular structure characterized by the enclosure of one or more cells into the cytosolic vacuoles of another cell. CICs had been mostly documented in various of human tumors^1,2^, such as pancreatic cancer ^3–5^, head and neck squamous cell carcinoma ^6^ and breast cancer ^7,8^, where it was proposed to mediate competition between tumor cells to facilitate clonal selection and tumor evolution ^9^. Meanwhile, recent studies implicated CICs in a wider range of biological processes, such as embryonic development ^10^, genome stability ^11,12^, virus infection ^13,14^ and immune homeostasis ^15,16^ and the like. Over the past decade, intensive efforts were endeavored to decipher the molecular mechanisms underlying the formation of CICs, among which entosis represents one of the mostly studied programs. During entotic CIC formation, the polarized adherens junction and contractile actomyosin are two well-established core elements ^17,18^. Recently, the vinculin-enriched mechanical ring was identified as a novel core element interfacing between adherens junction and actomyosin to coordinate cell internalization ^19^. Besides, a group of factors, by acting on the core elements, were identified to regulate CIC formation, such as PCDH7, CDKN2A, MTRF and LPA and the like ^20–25^.

Despite of the great progress made on the formation mechanisms for CICs, much less is known about the fate control of the internalized cell in CICs. It was proposed that entosis was a CICs-mediated type IV programmed death ^26^, which resulted in lysosomal entotic death as a predominant mechanism and apoptotic death as an alternate program when lysosomal function was disrupted ^27^. The LC3 lipidation onto the entotic vacuole seemed to be associated with the death of the internalized cells via facilitating its fusion with adjacent small lysosomes ^28^. The dead inner cells were cleared by entotic vacuoles, as a huge lysosome actually, through active fusion and fission processes ^29,30^. Nevertheless, previous studies of the death mechanisms of the internalized cells had not dealt with the critical condition of lysosome acidification for cell death and degradation, and the determined turning point for the fates of internalized cells.

We here introduced keima ^31,32^, a protein pH sensor, to measure the precise pH of entotic vacuoles during the death of internalized cells and explore the complementary relationship between entotic death and apoptotic death. Keima is a coral-derived acid-stable fluorescent protein that has a bimodal excitation spectrum peaking at 440 and 550 nm corresponding to the neutral and ionized states ^32^. Generally, the dual-excitation ratio (550 nm / 440 nm) of this probe can precisely indicate the cellular pH value, allowing its extensive usage in various studies of cell physiology. Keima was adopted to detect autophagic events based on lysosomal delivery ^33^, and was also applied as a marker of mitophagy ^34^ to explore autophagosome-lysosome fusion ^35^. With this genetically encoded fluorescent protein, we could conveniently probe the pH dynamics of cell death occurring within cell-in-cell structures.

In this study, we studied the pH titration behaviors of keima-overexpressed cells using gradient pH buffer, which enabled us to read out the context-specific pH value for each cell lines based on its correlation with the dual-excitation ratios of keima protein. The keima-overexpressed cells were then applied to the time-lapse assay to measure the accurate pH value of entotic vacuole during the process of the inner cell death associated with different patterns of LC3 lipidation. Moreover, we demonstrated that manipulating lysosomal acidification is effective in controlling inner cell fates as well as the ways whereby the inner cells die.

## Materials and methods

### Antibodies and chemical reagents

The following antibodies were used: anti-cleaved-caspase 3 (1:200; CST, #9664s) and secondary Alexa Fluor 647 anti-rabbit (1:500; Invitrogen, #A-20991). Reagents including EN6 (Selleck, #S6650), Hydroxychloroquine (CQ) (MCE, #HY-W031727), Concanamycin A (Con A) (Shanghai ZZbio.co, ZAE-ALX-380-034-C025), NH_4_Cl (Coolaber, CA30112320), LysoTracker (Invitrogen, #L7526), DAPI (Life technologies, #D1306) were purchased and used according to manufacturer’s instructions.

### Cells and culture conditions

Cell lines MCF7, SW480 expressing E-cadherin (SW480/E), MDA-MB-231 expressing E-cadherin (MM231/E), HEK293T, and their derivatives were cultured in Dulbecco’s modified Eagle’s medium (MACGENE Technology Ltd., Beijing, China) supplemented with 10% fetal bovine serum (Kangyuan Biology, China). MCF10A and their derivatives were maintained in DMEM/F12 (Gibco, USA) supplemented with 5% equine serum (Kangyuan Biology, China), 20 ng/ml EGF (Peprotech, USA), 10 g/ml insulin (Sigma, USA), 0.5 ug/ml hydrocortisone (Sigma, USA), and 100 ng/ml cholera toxin (Sigma, USA). All cells were cultured in the humidified incubator of 5% CO_2_ at 37°C.

### Cloning and generation of stable cell lines

pQCXIP-mKeima-N1 was cloned by inserting the mKeima sequence (pCHAC-mt-mKeima, Addgene, #72342) into *Bam*HI/*Sal*I sites of the pQCXIP-N1 vector using the T4 DNA ligase (New England BioLabs, #M0202S) according to manufacturer’s instructions. The cDNA encoding EGFP-LC3 fusion protein was released from pBABE-EGFP-LC3-puro (Addgene, #22405) and cloned into pQCXIN-EGFP-N1-Neo by the 5’-*Eco*RI and 3’-*Age*I sites. To generate stable cell lines, retroviruses were packaged in HEK293T cells using Lipofectamine 2000 reagent (Invitrogen) as described before ^36^. All cell lines were transduced with viruses for 24 h with 8 μg/ml polybrene (Sigma). Virus-infected cells were selected and grown in medium with 2 μg/ml puromycin or 200 μg /ml G418.

#### pH titration

Cells overexpressing fluorescent protein keima were incubated in a series of buffers with pH values ranging from 4.12 to 7.97, then the fluorescent signal of keima was measured under the conditions of excitation of 440 nm and 550nm and emission of 610 nm using Nikon ECLIPSE Ti-U epi-fluorescence microscope. Then the fluorescent intensity of keima was measured by NIS-Elements F 3.0 software ^37^. The pH titration curves and fitting equation were obtained by the negative correlation between the pH value and the fluorescent intensity ratio of 550 nm / 440 nm of keima protein.

### Time-lapse microscopy

Cell-in-cell time-lapse assay was performed as previously described ^2^. About 3 x 10^5^ cells were suspended in 0.5% agarose-coated plates for 6 hours, and then cell suspensions were transferred and grown on cover-glass dishes. Images of cells were captured for DIC and fluorescence channels (excitation of 440 nm and 550nm and emission of 610 nm) every 10 min with 20 x objective lenses at 37°C and 5% CO_2_ for 24 hours by the Nikon ECLIPSE Ti-U epi-fluorescence microscope and analyzed by NIS-Elements F 3.0 software (Nikon, Japan). The timing of cell death was judged morphologically by the appearance of a broken cell membrane, or cessation of cell movement, or both.

### LysoTracker staining

LysoTracker staining was performed according to manufacturer’s instructions. LysoTracker FITC (Invitrogen, #L7526) dissolved in serum-free medium was added to keima-expressing cells for 30 min at 37°C. After that, cells were washed with PBS and cultured in complete growth medium followed by imaging using Nikon ECLIPSE Ti-U epi-fluorescence microscope.

### Immunostaining

Immunostaining was performed as previously described ^38^. Briefly, cells were fixed in 4% paraformaldehyde for 10 min at room temperature, then permeabilized with 0.2% Triton X-100/PBS for 3 min and washed with PBS followed by blocking with 5% BSA at room temperature for 1 hour. Fixed samples were incubated with primary antibody at 4°C overnight and washed with PBS before incubated with fluorophore-labeled secondary antibodies at room temperature for 1 hour. Cells mounted with mounting medium with DAPI (ZSGB-BIO, #ZLI-9557) and imaged by Nikon ECLIPSE Ti-U epi-fluorescence microscope.

### Statistical analysis

All of the experiments were performed for at least three times. Data were displayed as mean ± SD. *P* values were calculated by two-tailed Student’s t test or Dunnett-t test using Excel or GraphPad Prism software, with statistical significance assumed at *P* < 0.05.

## Results

### Keima-based monitoring of the cellular and lysosomal pH values

To monitor the pH dynamics in live cells, we made cell lines that stably expressed keima, a genetically engineered protein pH meter that displays different excitation spectra upon pH changes. As shown in **Fig. 1A**, the excitation at 550 nm decrease accompanied with an increase in excitation at 440 nm when the buffering pH is getting higher, which is consistent with the published study ^33^. Briefly, keima displayed red in the acidified buffer (pH 4.12-5.5), green in the neutral buffer (pH 7.07-7.97), and orange in the transitory buffer (pH 5.6-7.06). For accurate pH measurement based on keima excitation, we managed to make a correlation between the pH value and the ratio of keima excitations (550 nm / 440 nm), and obtained pH titration curves for four entosis-proficient cell lines: MCF7, MCF10A, SW480 expressing E-cadherin (SW480/E) and MDA-MB-231 expressing E-cadherin (MM231/E), respectively (**Fig. 1B-E**). Interestingly, based on the titration curves and the derived equations, the above four cell lines were found to have profoundly different intracellular pH values (**Fig. S1B, C**).

**Fig. 1.**
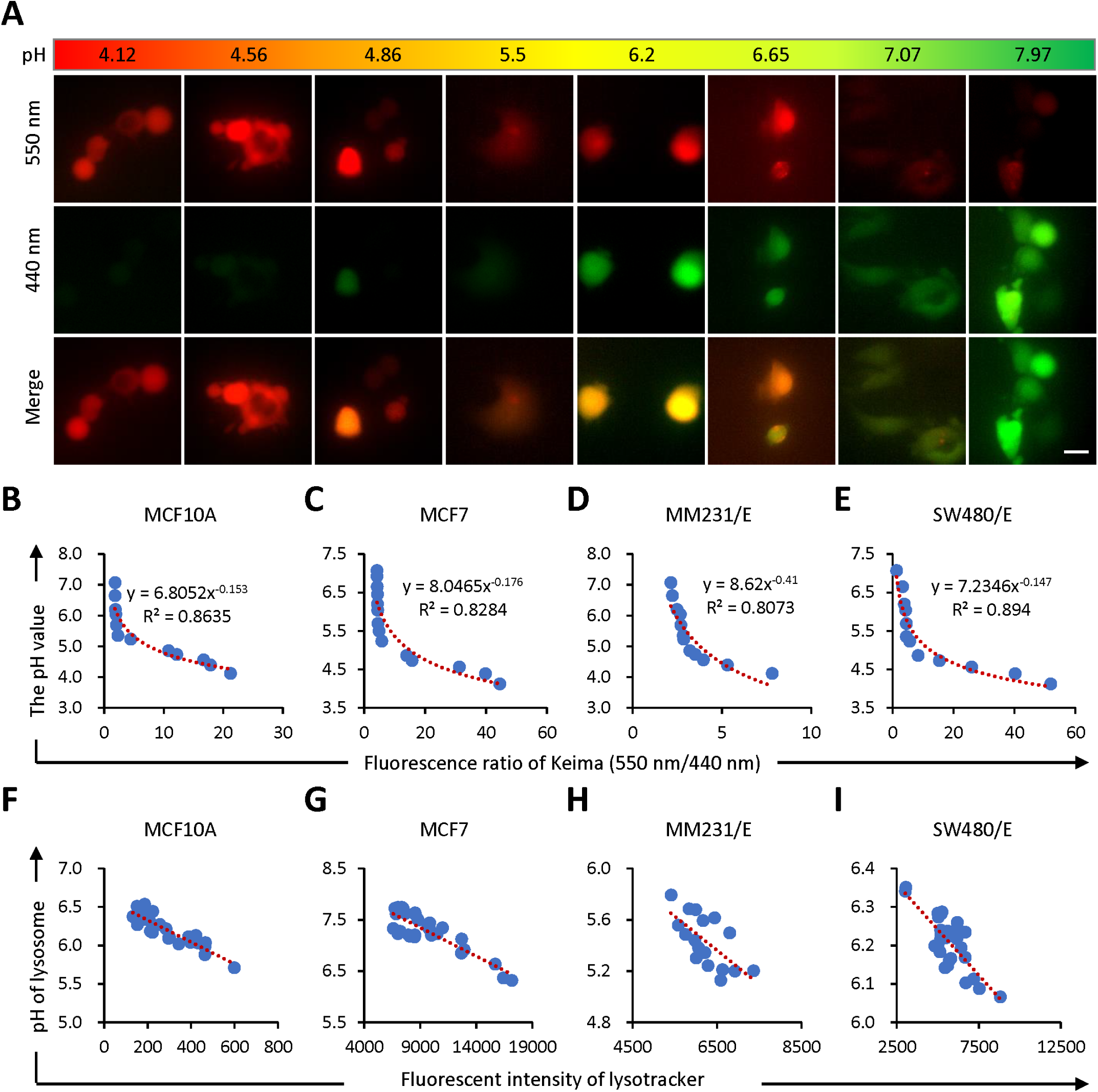
Probing cellular pH by keima fluorescent protein. **(A)** Representative images of three channels (550 nm, 440 nm and merged) for keima-overexpressed MCF7 cells in a series of buffers of different pHs. Scale bar, 20 μm. **(B-E)** The cellular pH value was negatively correlated with the excitation ratio of keima (550/440), in MCF10A (B), MCF7 (C), MDA-MB-231/E (D) and SW480/E (E), respectively. **(F-I)** The lysosomal pH was negatively correlated with the fluorescent intensity of LysoTracker in MCF10A (F), MCF7 (G), MDA-MB-231/E (H) and SW480/E (I), respectively.

In agreement with its pH sensitive property, keima displayed bright red fluorescent signals (550 nm) at the subcellular regions marked by lysotracker, a fluorescent dye specifically labeling lysosomes (**Fig. S1A**) ^39^. Remarkably, the pH values derived from the keima ratio of 550 nm / 440 nm were tightly correlated with the intensity of lysotracker at the subcellular organelle of lysosomes (**Fig. 1F-I**), further supporting the liability of keima to be an *in situ* pH meter for live cell analysis.

### The pH dynamics during the death of internalized cells

By taking advantage of the keima-expressing cells as a pH read out, we firstly tried to examine the pH changes of the entotic vacuoles during the inner cell death of MCF10A cells. As shown in **Fig. 2A**, the entotic vacuoles gradually turned into yellow and subsequently into red along with the death and degradation of the inner cells, indicating progressive vacuolar acidification (**Fig. S3A and Movie S1**). Based on the changes in keima excitation and inner cell morphology, the death process could be roughly divided into four steps, including internalization (S1), acidification (S2), entotic death (S3), and degradation (S4) (**Fig. 2B**). While the pH changed little in the outer cells, it kept declining in the dying inner cells during the four sequential stages (S1-S4) (**Fig. 2B, C**). By contrast, the live inner cells were in green throughout whole process of time lapse imaging (**Fig. S3B and Movie S2**), which is similar to those of the outer cells and non-CICs live cells (**Fig. S3C**), indicating relative stable pH of the entotic vacuoles containing live cells.

**Fig. 2.**
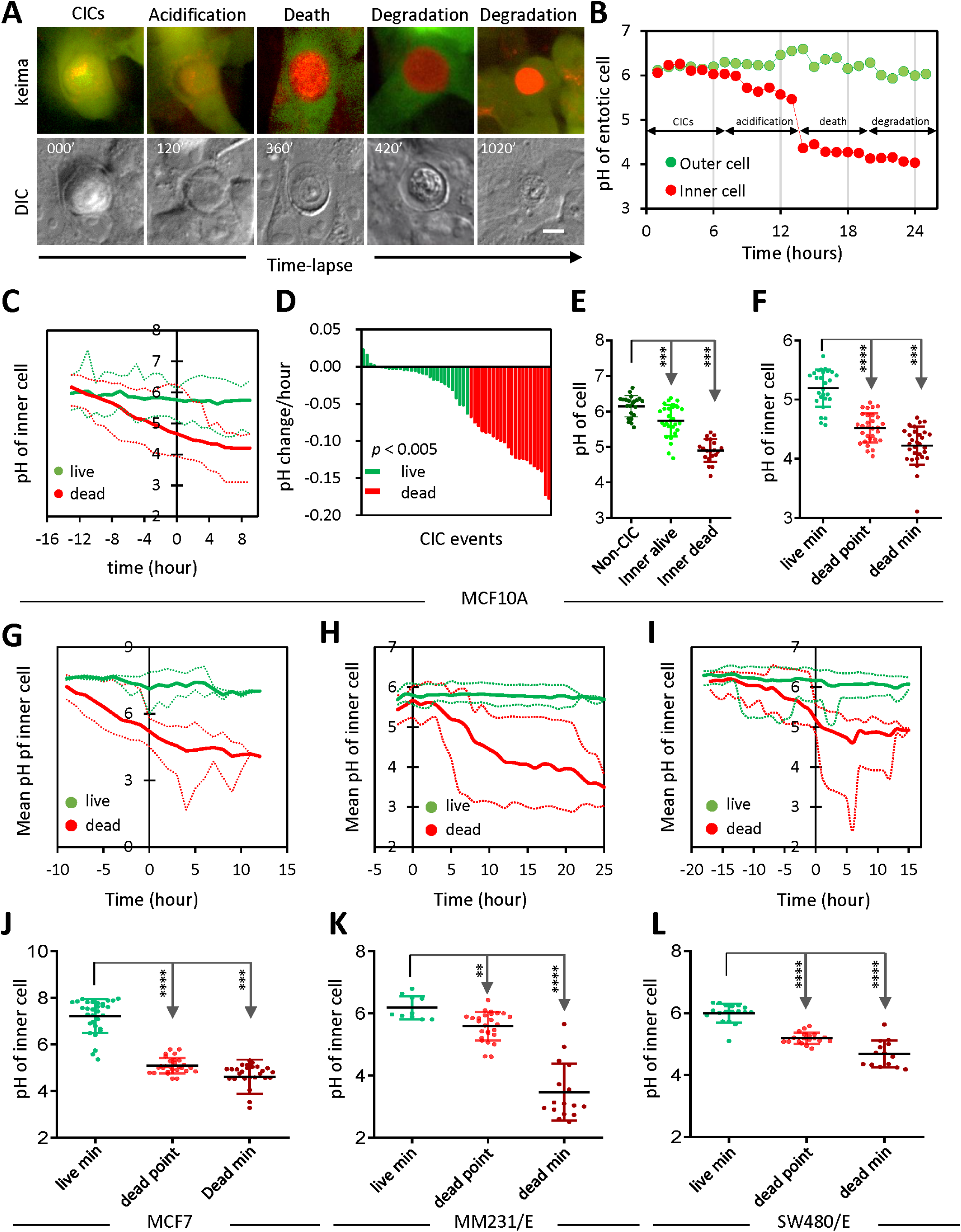
The pH dynamics during the CICs-mediated death. **(A)** The time-lapse images showed keima signal changes in one CICs from MCF10A cells. The keima channel was merged from 440 nm (green) and 550 nm (red) channels. Scale bar, 10 μm. **(B)** The quantitative presentation of pH changes in (A) for the respective inner and outer cell over time course. **(C)** The pH changes of inner cells from a group of CICs over time. The solid lines are for the mean pH while the upper dotted lines and lower dotted lines indicate the pH ranges. The inner cell death was set as time zero. N = 17 (dead), 28 (live). **(D)** The change of pH value per hour for live (green) and dead inner cells (red), respectively. **(E)** Quantification of the cellular pH for non-CIC cells (dark green), live inner cells (green dots) and dead inner cell (dark red). **(F)** Graph shows the minimum pH of live inner cell (green), the pH of dead inner cells at the moment of cell death (bright red) and minimum pH of the dead inner cell (dark red). **(G-I)** The pH of inner cells changed over time in the keima-overexpressed MCF7 (G), MDA-MB-231/E (H) and SW480/E (I), respectively. (G) n = 12 (dead), 8 (live); (H) n=12 (dead), 17 (live); (I) n=19 (dead), 18 (live). **(J-L)** Graph shows the minimum pH of live inner cell (green), the pH of dead inner cells at the moment of cell death (bright red) and minimum pH of the dead inner cell (dark red) for MCF7 (G), MDA-MB-231/E (H) and SW480/E (L), respectively. ** *P*<0.01; *** *P*<0.001; **** *P*<0.0001.

For the sake of more accurate evaluation, we calculated the rate of the pH changes in different categories of cells, and found that the pH of vacuoles with dying cells decreased at a speed of 0.0678 to 0.1779 per hour that was significantly faster than those for vacuoles with live cells (−0.0236 to 0.0632 per hour) (**Fig. 2D**). The mean pH of the non-CIC normal cells (6.14) was slightly higher than that of the alive inner cells (5.74) which was significantly higher than that of dead inner cells (4.90) (**Fig 2E**). Importantly, the pH of 4.5 seemed to be the dead point, referring to dead pH hereafter, for the internalized MCF10A cells as no cells were found to be alive at pH below 4.5, whereas, cells would be alive within vacuoles of pH > 5.0 (**Fig. 2F**). Similar dynamics were obtained on the other three entosis-proficient cells (**Fig. 2G-I**). However, the dead pH varied with 5.0 for MCF7 cells (**Fig. 2J**), 5.6 for MDA-MB-231/E cells (**Fig. 2K and S3D, E**), and 5.2 for SW480/E cells (**Fig. 2L and S3F, G**). Interestingly, the dead pHs for the later three cells that are cancer cells were much higher than that of MCF10A cells that are non-transformed epithelial cells (**Fig. 2F, 2J-L**). These data suggest that the vacuolar pH is critical for the fate control of the internalized cells, which, however, is context-dependent, and cancer cells seemed to be less tolerant to the acidified environments than the non-cancerous epithelial cells.

### LC3 lipidation onto the entotic vacuoles in complex patterns

LC3 lipidation of the entotic vacuole was reported to be a critical event taking place right prior to and mediating entotic cell death by facilitating lysosome fusion ^28^. It is interesting to determine the relationship between LC3 lipidation and acidification of the entotic vacuoles. We therefore established MCF10A cells stably co-expressing keima and EGFP-LC3 and examined the LC3 recruitment to the entotic vacuoles by time-lapse microscopy of 24 hours as the first step. As reported, the typical LC3 lipidation, which is rapid, transient and generally occurred within one hour, of the entotic vacuole were observed right before the entotic death in most of the CICs (53.3%) (**Fig. 3A-C and Movie S3**). Unexpectedly, a number of vacuoles (40%) also recruited LC3 post inner cell death, which could be subdivided into two classes based on the presence of pre-death LC3 lipidation, ie, LC3-entosis-LC3 (**Fig. 3D**) and entosis-LC3 (**Fig. 3E and Movie. S5**). Moreover, we observed two cases of LC3 lipidation that were not associated with any signs of inner cell death in the 24 hours duration of time lapse (**Fig. 3F and Movie S6**), instead, LC3 either retained for long time (200 min), or was repetitively recruited for multiple times. Furthermore, the post-death LC3 lipidation seemed to be an indispensable event (100%) for the vacuoles that contained cells dying through an apoptotic program (**Fig. 3G, H and Movie S4**). These finding demonstrated a previously underappreciated relationship between LC3 lipidation and cell death mediated by cell-in-cell structures.

**Fig. 3.**
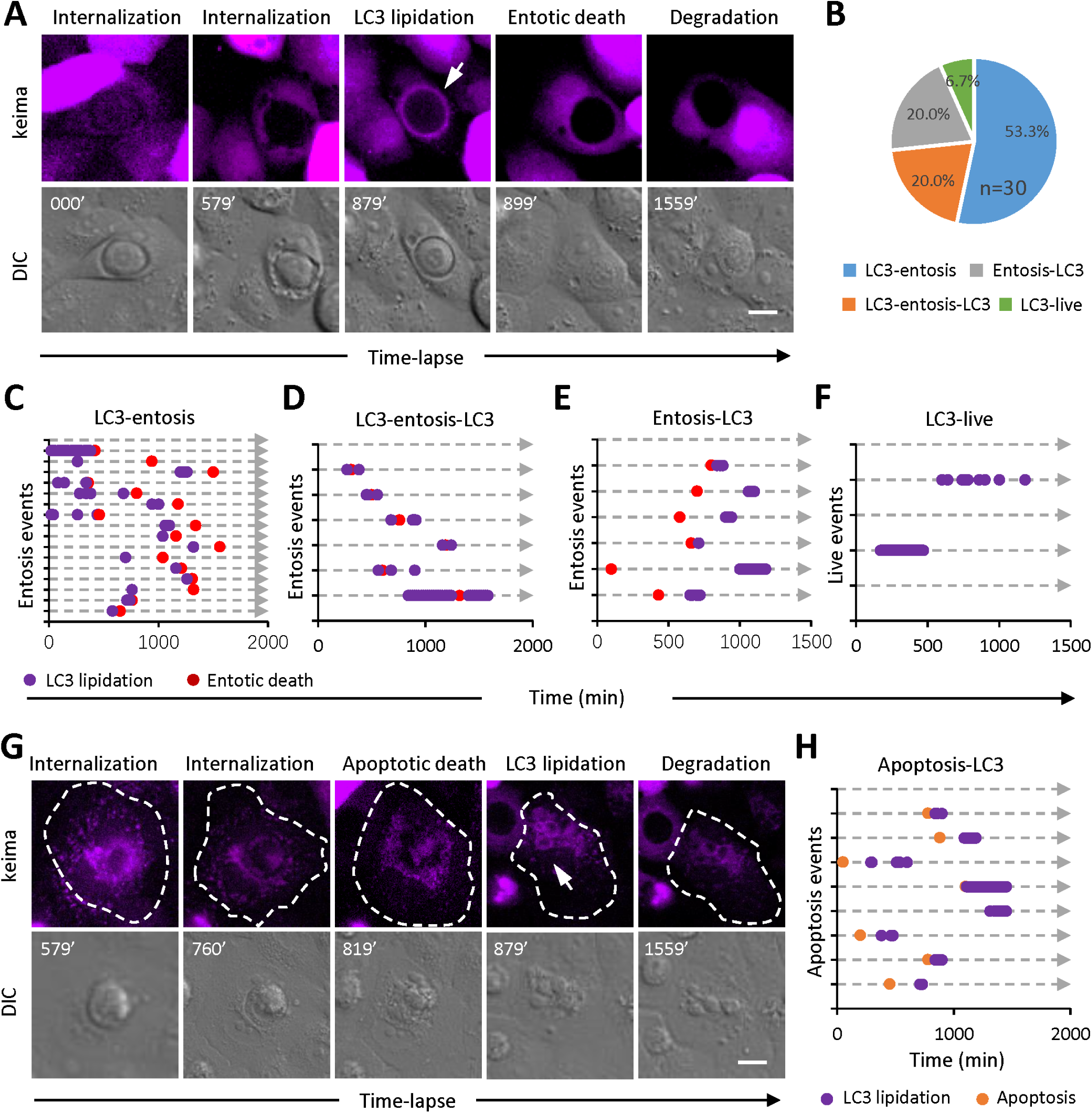
The dynamics of vacuolar LC3 lipidation. **(A)** Time-lapse images of transient recruitment of GFP-LC3 (purple) to the entotic vacuole prior to inner cell death. Arrow indicates LC3 on the entotic vacuole, also see Supplementary Information Movie S3. Scale bar, 10 µm. **(B)** Pie chart shows the percentage of four types of LC3 lipidation as indicated, n = 30. **(C-F)** Time course-based plotting of four patterns of LC3 recruitment (purple dots) in related with the fates of internalized cells, including LC3 lipidation before entotic death (C), LC3 lipidation before and after entotic death (D), LC3 lipidation after entotic death (E), LC3 lipidation for live inner cells (F). **(G)** Time-lapse images of vacuolar GFP-LC3 (purple) recruitment following inner cell apoptosis. Arrow indicates LC3 recruited to the apoptotic body, also see Supplementary Information Movie S4. Scale bar, 10 µm. **(H)** Time course-based plotting of LC3 lipidation in related with inner cell apoptosis.

### LC3 lipidation associated with vacuolar acidification during the death of internalized cells

We next investigated the pH change associated with the LC3 lipidation with the co-expressed keima protein. All the pH values associated with the above five patterns of LC3 lipidation were measured *in situ* and plotted along the time course of death events (**Fig. 4C, D and S4A-C, E, F**). For the typical LC3 lipidation that occurred prior to the entotic death of inner cells (**Fig. 3A**), the pH of entotic vacuoles was in a range of 4.8-6.20 followed by the entotic death of the inner cells at a lower pH about 4.3-5.2 (**Fig. 4A, C and Movie S3**). By contrast, the apoptotic death of the inner cells took place at a relatively higher pH about 5.3-6.0 followed by LC3 lipidation on the apoptotic bodies at a lower pH about 4.5-5.6 (**Fig. 4B, D and Movie S4**). And the pH for LC3 lipidation of post-entotic death was about 4.4-4.8, which was frequently preceded by entotic death events of higher pH (4.8-5.6) as compared with that of entotic death without post-death LC3 lipidation **(Fig. 4E-G and S4A, B)**. The pH for LC3 lipidation without death was similar to that of pre-entotic death **(Fig. 4G and S4A, C)**. Nevertheless, all the LC3 lipidation events were followed by a continuous decline of vacuolar pH **(Fig. 4H)**, which is consistent with a role of LC3 lipidation in facilitating entotic vacuoles-lysosome fusion ^28,40^. These data suggested that LC3 lipidation may not occur in responding to a defined range of vacuolar pH, instead, vacuolar acidification was a consequence of LC3 lipidation that not only initiated entotic cell death but also facilitated subsequent degradation of corpses from both entotic and apoptotic death. Consistently, LC3 lipidation took place primarily in entotic vacuoles containing dead cells, either entotic (82.4%) or apoptotic (80.0%), but rather infrequent in vacuoles with live cells (2.4%) (**Fig. S4D**).

**Fig. 4.**
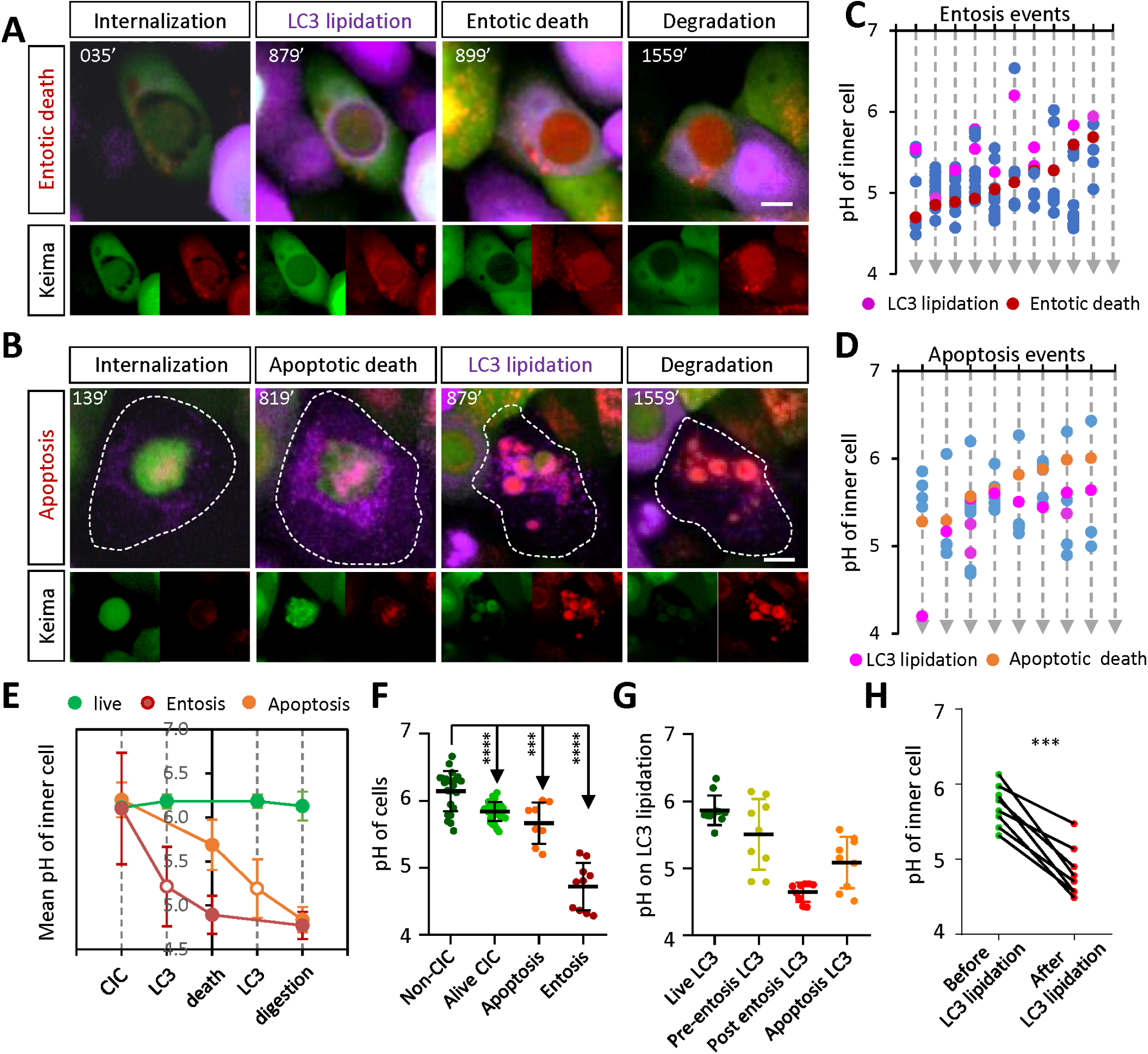
The analysis of vacuolar acidification along with LC3 lipidation. **(A, B)** Time-lapse images for keima (excitation at 440 nm in green, 550 nm in red) and GFP-LC3 (purple) during the entotic (A), or apoptotic (B) death of internalized MCF10A cell. Scale bars, 10 μm. **(C, D)** Time course-based plotting of vacuolar pH in related to LC3 lipidation during the entotic (C), or apoptotic (D) death of internalized cells. **(E)** The average pH dynamics for inner cells that were alive (green), or underwent apoptotic death (orange), or underwent entotic death (red), respectively. **(F)** The pH for normal cells (non-CICs), alive inner cells, apoptotically dead inner cells and entotically dead inner cells, respectively. **(G)** The vacuolar pH on the moment of LC3 lipidation, including LC3 lipidation for alive inner cells (green), LC3 lipidation before entosis (yellow), LC3 lipidation after entosis (red) and LC3 lipidation after apoptosis (orange). **(H)** Changes in vacuolar pH before and after LC3 lipidation. ** *P*<0.01; *** *P*<0.001; **** *P*<0.0001.

### The fate control of internalized cells by vacuolar pH manipulation

Given the pivotal role of pH in determining inner cell death, it’s conceivable that manipulating vacuolar acidification might be a way to control inner cell fates. To examine this idea, we first treat MCF10A cells with EN6, an activator of v-ATPase that facilitates vacuolar acidification ^41^, which significantly lowed the pH of entotic vacuoles containing live cells to around 5.1, a value really close to the range of the death pH (4.0-4.9) (**Fig. 2F and 5A**). As a result, the death rate of inner cells increased (from ∼45% to >65%) (**Fig. 5B**), and, remarkably, all the death events were exclusively entotic (100%) but not apoptotic (**Fig. 5C**), as confirmed by immunostaining of cleaved-caspase 3, the marker of apoptosis (**Fig. 5D, 5E**). To our best knowledge, this is the first demonstration that enhancing vacuolar acidification could promote CICs-mediated entotic death of internalized cells.

**Fig. 5.**
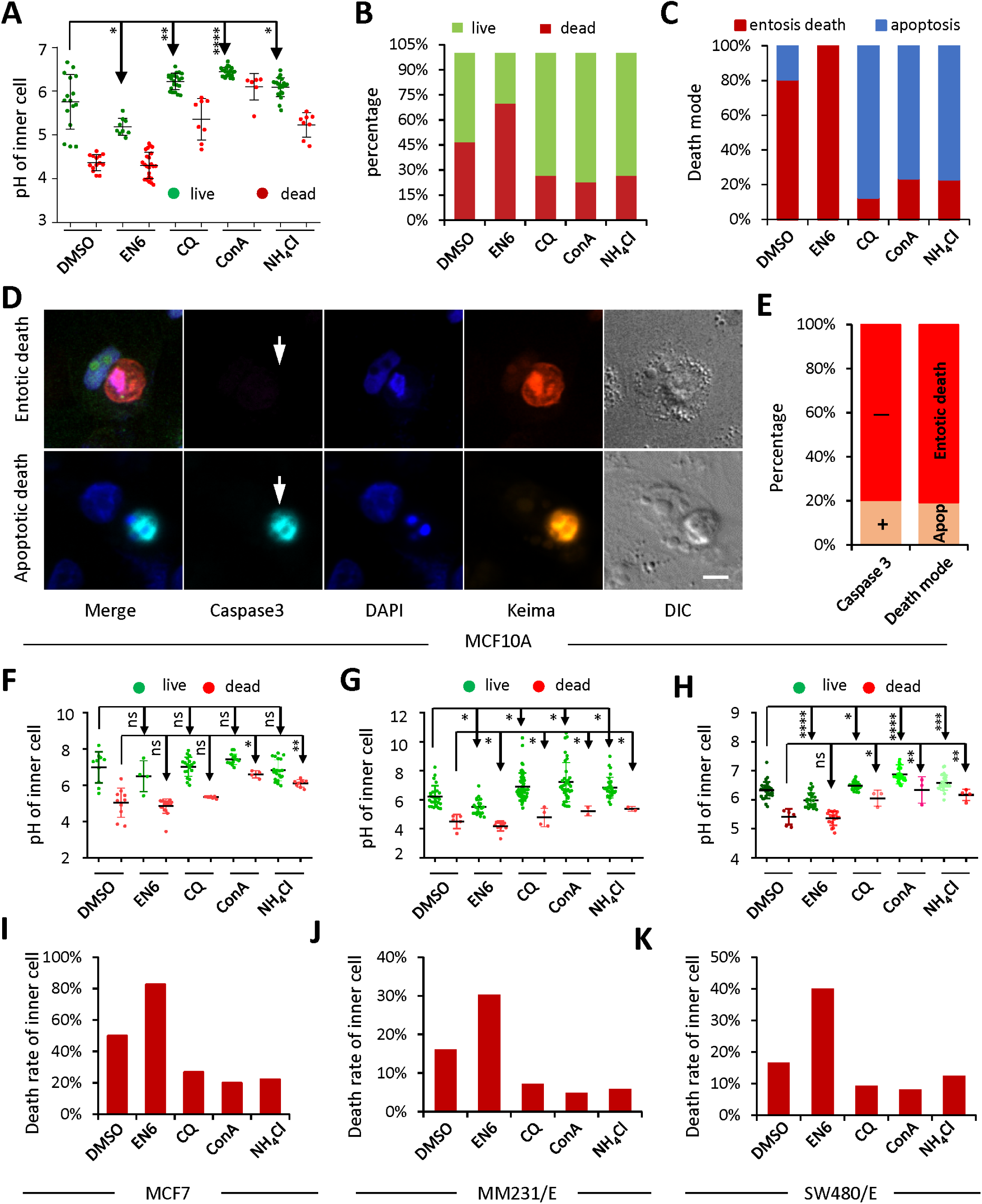
The fates of inner cells were controlled by vacuolar pH. **(A-C)** The pH of vacuoles containing live or dead inner cells (A), and inner cell fates (B), and the ways inner cells died (entotic or apoptotic) (C), for MCF10A cells upon treatments with EN6 (10 μM), CQ (20 μM), ConA (200 nM) and NH_4_Cl (100 μM). n (left to right) = 30, 33, 30, 31, 30 for (A); 30, 33, 30, 31, 30 for (B); 58, 49, 32, 34, 26 for (C). **(D)** Representative images of CICs, with inner cells undergoing entotic death (up) or apoptotic death (low), that were stained with antibody for cleaved-caspase 3 (cyan). Keima for merged channels of 440 nm and 550 nm. White arrows indicate inner cells. Scale bar, 10 μm. **(E)** Quantification of inner cells that were positive in cleaved-caspase 3, or died either entotically or apoptotically. n (left to right) = 64, 57, respectively. **(F-K)** The pH of vacuoles containing live or dead inner cells (F-H), and inner cell fates (I-K), for cells (MCF7, MM231/E and SW480/E) upon treatments with EN6 (10 μM), CQ (20 μM), ConA (200 nM) and NH_4_Cl (100 μM), respectively. n (left to right) = 26, 23, 26, 20, 28 for (F, I); 31, 32, 57, 38, 33 for (G, J); 36, 43, 32, 37, 32 for (H, K). * *P*<0.05; ** *P*<0.01; \*\*\**P*<0.001; **** *P*<0.0001.

To confirm this finding, we then treated MCF10A cells with compounds that were capable of compromising vacuolar acidification, including concanamycin A (ConA) that selectively inhibits V-ATPase ^42^, chloroquine (CQ) that could inhibit vacuole-lysosome fusion ^43^, and ammonium chloride (NH_4_Cl) that could alkalinize intracellular compartments ^44,45^. As expected, the average pH of the entotic vacuoles containing live cells increased to above 6.1 upon CQ, or ConA, or NH_4_Cl treatments (**Fig. 5A**), leading to much less inner cell death (<30%) as compare with control (>45%) and EN6-treated cells (>65%) (**Fig. 5B**), and, interestingly, majority of the death were switched to apoptosis (from ∼20% to ∼80%), and consistent with above observation (**Fig. 4C**), the mean death pH for apoptosis here is of >5.2 (**Fig. 5A**). Furthermore, similar results were obtained from the other three CICs-proficient cells, including MCF7 (**Fig. 5F, I, S5**), MDA-MB-231 (**Fig. 5G, J**) and SW480 cells (**Fig. 5H, K**), and among them, the mean pH of each case was generally higher than that in MCF10A cells (**Fig. 5A, F-H**). Together, these results support that the inner cell death of a CIC structure was dictated by the vacuolar pH and could be manipulated by compounds that regulate the acidification of intracellular vacuoles.

## Discussion

We studied the dynamic process of pH change in the internalized cells in the CICs by keima, a fluorescent protein pH meter. We determined the lysosome-dependent death pH critical for the entotic death (**Fig. 6**), and identified LC3 lipidation onto the entotic vacuole as a preceding condition for the pH declining to a critical value and evoke the entotic death. Meanwhile, the LC3 lipidation following cell death, primarily apoptotic and, to a lesser extends, entotic, would promote the clearance of dead corpse by facilitating vacuolar acidification.

**Fig. 6.**
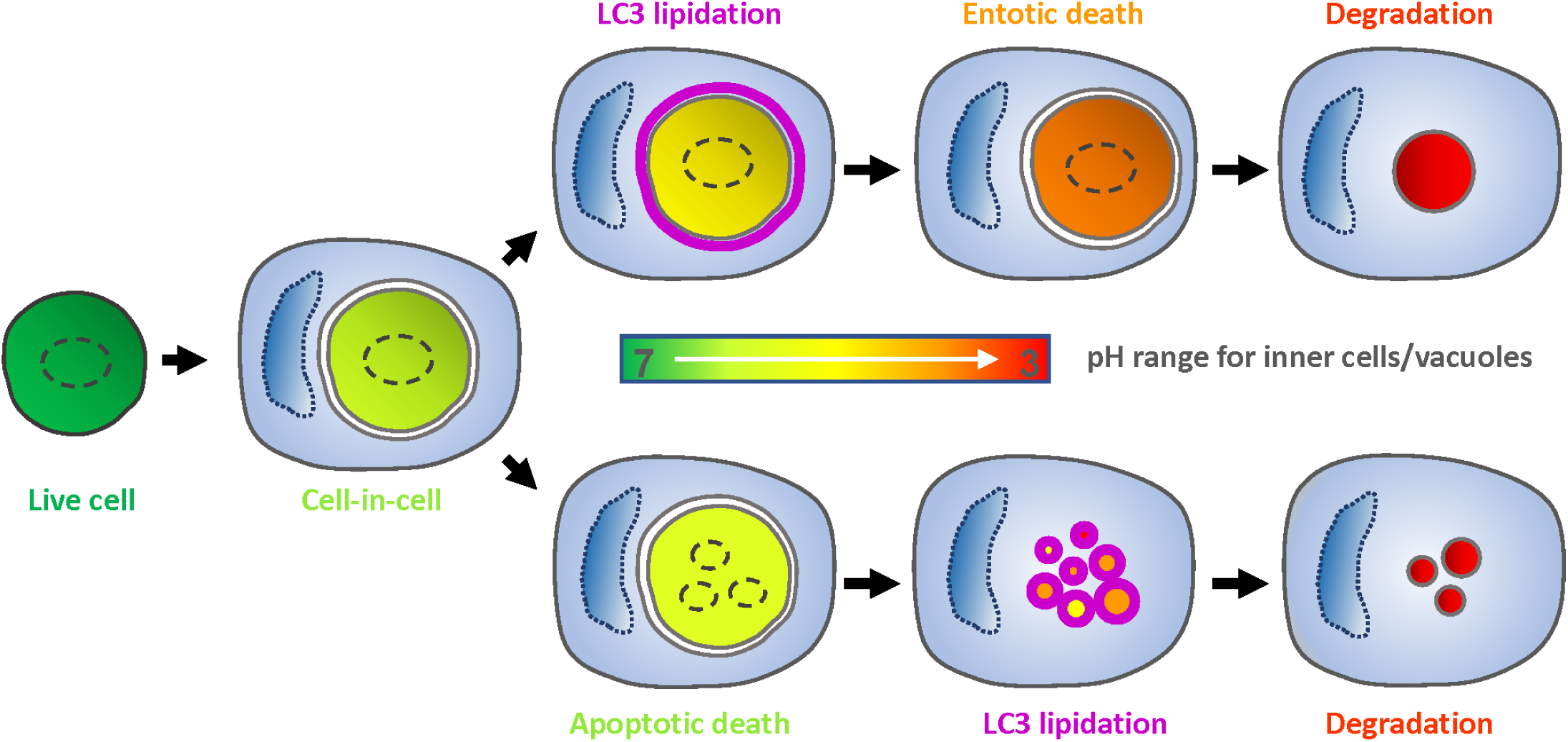
Schematic demonstration of pH dynamics along with typical LC3 lipidation in two ways of inner cell death (entotic or apoptotic) mediated by CICs. Following the fully internalization to form CICs, the inner cells generally die via two mechanistically distinct ways. On one hand, inner cells could be killed non-autonomously by the outer cells that are lysosome-dependent and preceded by a transient pre-death LC3 lipidation onto the entotic vacuole; on the other hand, inner cells may die cell-autonomously in a caspase-dependent apoptotic way, which is followed by a rapid post-death LC3 lipidation onto the apoptotic vacuoles. Under both circumstances, the pH of vacuoles that contain the internalized cells continues to decline, which may function to kill the inner cells and/or facilitate the clearance of dead corpses. Colors ranging from green to red indicate pH rang from neutral (about 7) to acidity (about 3). Purple ring indicates LC3 lipidation to vacuole.

It is known that fluorescent dyes, such as Lysotracker, could efficiently label acidified intracellular vacuoles and emit bright signals, and were generally employed to detect acidified vacuoles ^39^. However, these dyes, due to strong phototoxicity and easily to be quenched, were not good candidates for long period of time lapse imaging, particular for the process of CIC-mediated death that usually takes place within a period of up to >10 hours. Whereas, the fluorescent proteins, such as keima, were ideal alternatives attributed to their properties of optional excitation spectra and ratio-based measurement ^46^. As a proof of concept for the application in CIC-mediated death, we chosen four entosis-proficient cell lines, and examined the feasibility of keima as a pH meter to probe the role and dynamics of vacuolar acidification along with the inner cell death by long period (∼24 hours) of time lapse microscopy, which reported ideal results. Meanwhile, our study uncovered a number of previously unrecognized findings in terms of pH changes during CIC-mediated death, which set a basis for further investigation.

It is worth to pointed out that the pH of cells was correlated to the pH of lysosome in these four cell lines. The cytoplasmic pH and lysosomal pH of the MCF7 cells were the highest whereas those of the MDA-MB-231 cells were the lowest (**Fig. S1B, C**). Moreover, we found that lysosome pH and cellular pH were both positively related to entotic death rate for these four cell lines (**Fig. S1B-D**), suggesting that the cellular or even environmental pH might have profound impacts on cells’ behaviors. Indeed, there were studies showing that cancer cells were actually grown in acidic tumor microenvironment ^47,48^, under which circumstance, MDA-MB-231 cells displayed high metastatic potentials than did MCF7 cells ^49^, consistent with the idea that the cytoplasmic pH and lysosomal pH were somehow related to the metastasis and malignancy of the tumor cells.

Interestingly, in addition to a clear role of vacuolar pH in dictating inner cell fates and the ways whereby inner cells die, complicated relationships were observed among the pH titration behaviors of the four cell lines (**Fig. 1B-E**). For instances, the cellular pH and the lysosomal pH were positively correlated to entotic death time for MCF10A, SW480, MDA-MB-231 cell lines but not for MCF7 cell line (**Fig. S1B, C, S2C**); the rate of pH change was correlated to the death rate of the inner cells for MCF7, MCF10A, SM480 cell lines but not for MDA-MB-231 cell line (**Fig. S1D, S2 A**); despite of the negative correlation between the rate of pH change and the death time of the inner cells (**Fig. S2D-G**), there was no correlation between the death pH and the lysosome pH (**Fig. S1B, S2B**). The above complicated relationships might be due to the significant genotypic and proteomic differences among these cell lines, or were related to the cell metabolism regulated by lysosome as reported ^50,51^.

As our results suggested that vacuolar pH levels were not a trigger, but a consequence instead, of LC3 lipidation onto the entotic vacuoles, and an interesting issue, though might be beyond the scope of this study, is what are the upstream factors that dictate the initiation of LC3 lipidation. Though the LC3 lipidation was executed in the outer cells, the commanding signals were likely from the internalized cells as the apoptosis of inner cells seemed to be always followed shortly by the LC3 lipidation. Actually, previous study indicated that osmotic changes in the vacuoles could efficiently induce LC3 lipidation, which required activity of the vacuolar-type H (+)-ATPase (V-ATPase) ^42^. Thus, it’s conceivable that LC3 lipidation followed the entotic death might be a result of increased osmotic changes in the vacuoles, which ended up with increased vacuolar pH that facilitates the degradation and clearance of the dead corpse. Since CIC structures in tumor cells were believed to promote cell competition, clonal selection and tumor evolution by multiple mechanisms, such as conferring growth advantages to the outer survivors via consuming the dead inner cells ^1,9^, interfering with the vacuolar acidification might be a potential strategy for the treatment of cancers undergoing active CIC formation.

## Acknowledgements

We thank Dr. Zhongyi Wang for comments on manuscript preparation. This work was supported by the National Key Research and Development Program of China (2019YFA09003801 and 2018YFA0900804), the National Natural Science Foundation of China (31970685, 31671432 and 81872314), the Fundamental Research Funds for the Central Universities (WK2090050042) and Anhui Province Science and Technology Major Project (201903a07020019).

## Author Contributions

Concept and design: QS and TL; Phenotype: YS; Gene cloning: HR and BZ; Data collection: YS, HR, XH, ZN and LG; Figures: YS, QS, TL, MT and YZ; Data interpretation: QS, TL and YS; Manuscript: QS, TL and YS, with input from MT, YZ, ZN, CW and HS; Funding: QS, YZ, ZC and TL. All authors have read and approved the final manuscript.

## Supplemental Data

### Supplemental Figures

**Fig. S1.**
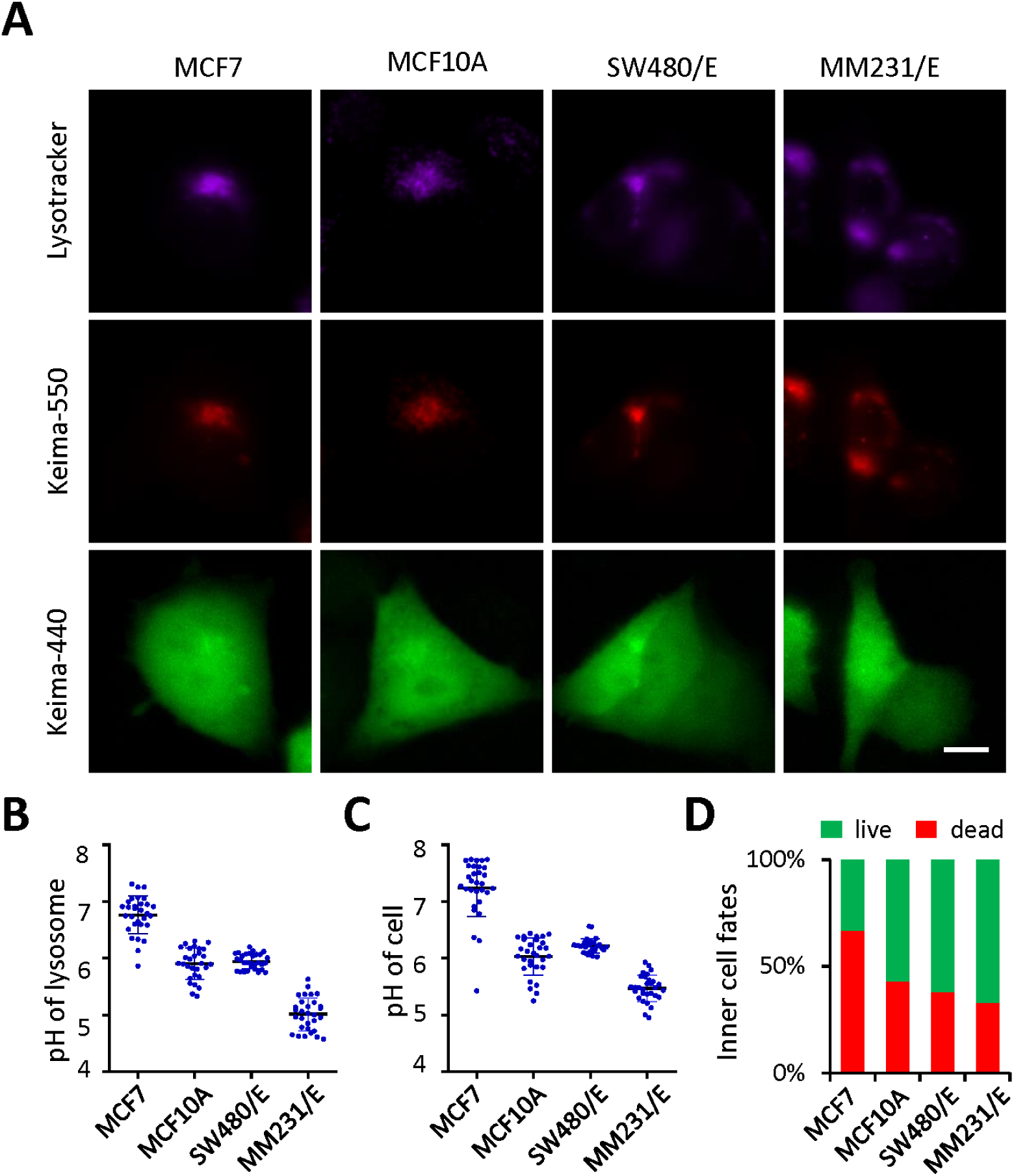
Keima-based measurement of the lysosomal pH and cellular pH. **(A)** Representative images for four keima overexpressed cell lines (550 nm, 440 nm), that were stained with LysoTracker (purple). Scale bar, 20 μm. **(B, C)** The lysosomal Ph (B) and cellular pH (C) for four keima overexpressed cell lines. **(D)** The ratio of inner cell fates (live/dead) for four cell lines.

**Fig. S2.**
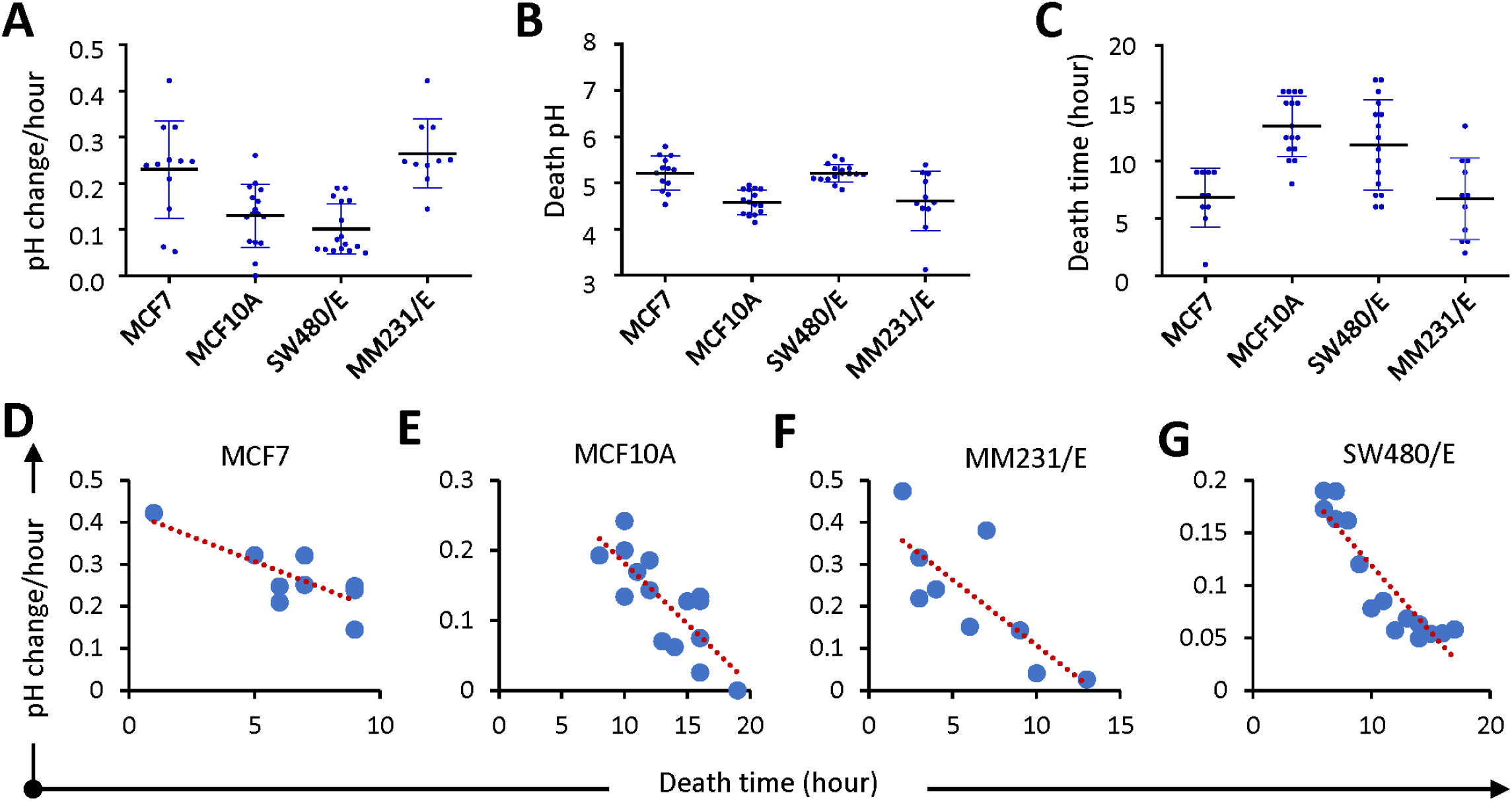
The change of pH value and the entotic death time. **(A-C)** The change of pH value per hour (A), the cellular pH on death moment (B) and death time (from internalization to cell death) (C) for dead inner cells in MCF7, MCF10A, MDA-MB-231/E and SW480 /E, respectively. **(D-G)** The rate of pH change was negatively correlated with the death time of inner cells in MCF7 (D), MCF10A (E), MDA-MB-231/E (F) and SW480 /E (G), respectively.

**Fig. S3.**
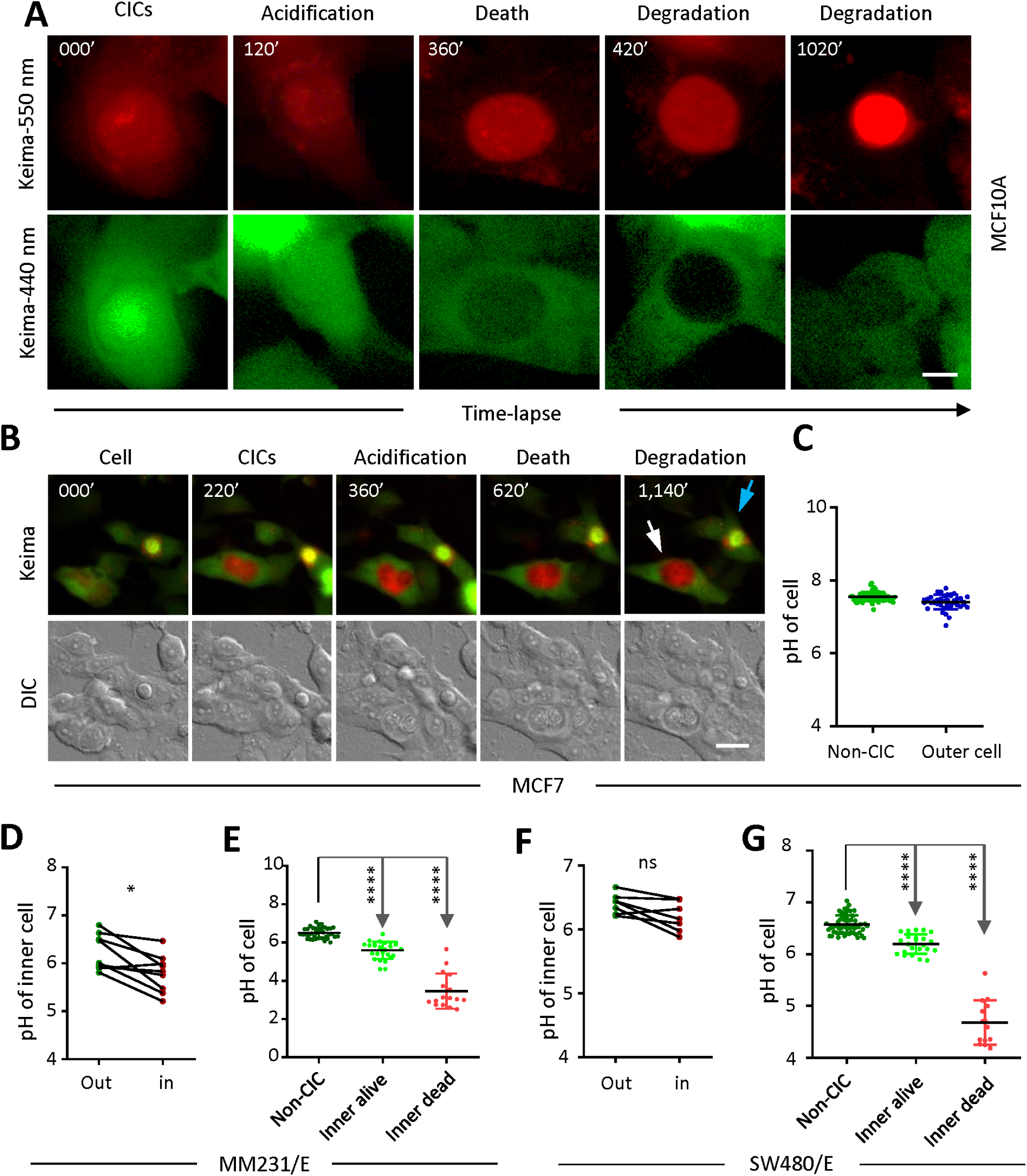
The pH dynamics during the CICs-mediated death. **(A)** The time-lapse images showed keima signal by two channels (550 nm and 440 nm) in one CICs from MCF10A cells. Scale bar, 10 μm. **(B)** The time-lapse images showed keima signal changes in two CICs from MCF7 cells. Blue and white arrows indicate live and dead inner cells respectively. Scale bar, 10 μm. **(C)** Quantification of the cellular pH for non-CIC cells (green) and outer cells of CICs (blue). **(D, F)** Changes in vacuolar pH before and after internalization. **(E, G)** Quantification of the cellular pH for non-CIC cells (dark green), live inner cells (green dots) and dead inner cell (red).

**Fig. S4.**
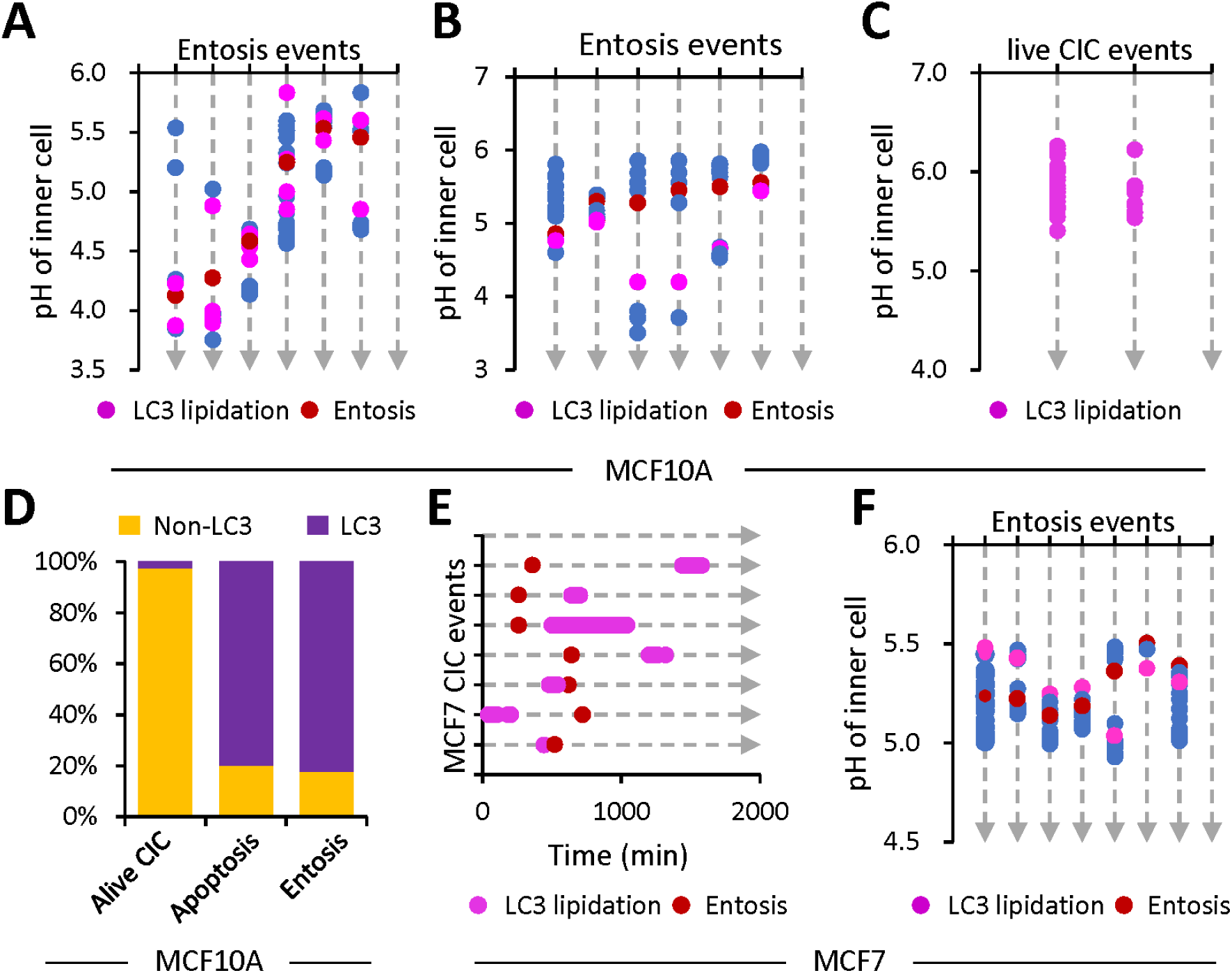
The analysis of vacuolar acidification along with LC3 lipidation. **(A-C)** Time course-based plotting of vacuolar pH in related to LC3 lipidation during the entosis, including LC3 lipidation before and after entotic death (A), LC3 lipidation after entotic death (B) and LC3 lipidation for live inner cells (C). **(D)** The ratio of LC3 lipidation in three situations of inner cell fate. n (left to right) = 81, 10, 34, respectively. **(E, F)** Time course-based plotting (E) and vacuolar pH (F) in related to LC3 lipidation during the entosis.

**Fig. S5.**
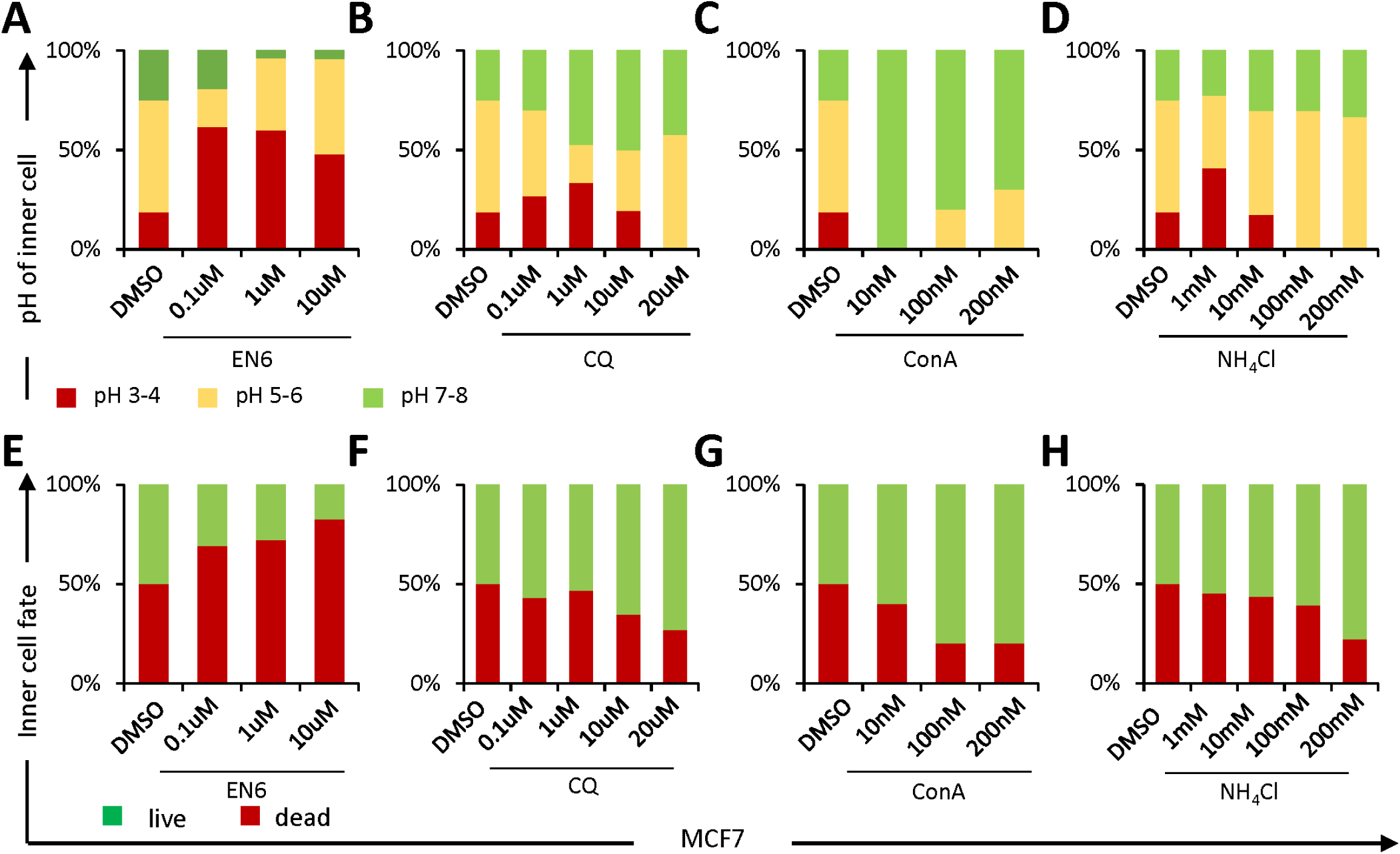
The vacuolar pH and cell fates of CICs treated with lysosome inhibitor. **(A-D)** The distribution of vacuolar pH upon treatments with different concentration of EN6 (A), NH_4_Cl (B), CQ (C) and ConA (D), respectively. The pH value was divided into three level: pH 3-4 (low, red), pH 5-6 (middle, yellow) and pH 7-8 (high, green). **(E-H)** The ratio of inner cell fates (live/dead) for MCF7 cells upon treatments with different concentration of EN6 (E), NH_4_Cl (F), CQ (G) and ConA (H) n (left to right) = 16, 25, 25, 23 for (A, E); 16, 30, 21, 26, 26 for (B, F); 16, 10, 20, 20 for (C, G); 16, 22, 23, 23, 18 for (D, H).

## Supplemental Movies

**Movie. S1** The time-lapse microscopy movies showed keima signal changes in one CICs from MCF10A cells.

**Movie. S2** The time-lapse microscopy movies showed keima signal changes in two CICs from MCF7 cells.

**Movie. S3** Time-lapse microscopy movies of transient recruitment of GFP-LC3 (purple) to the entotic vacuole prior to inner cell death.

**Movie. S4** Time-lapse microscopy movies of vacuolar GFP-LC3 (purple) recruitment following inner cell apoptosis.

**Movie. S5** Time-lapse microscopy movies of vacuolar GFP-LC3 (purple) recruitment following inner cell entosis.

**Movie. S6** Time-lapse microscopy movies of vacuolar GFP-LC3 (purple) recruitment for alive inner cells.

